# Biodesign Buddy: Integrating Generative Artificial Intelligence in Academic Biodesign

**DOI:** 10.64898/2026.03.11.710906

**Authors:** Dylan Riffle, Paul Rubery

## Abstract

Biodesign is an interdisciplinary research domain that incorporates principles from design and the life sciences to develop new systems, processes, and objects. Collegiate biodesign educators face unique pedagogical challenges, including an absence of relevant scholarship on curriculum design and instructional best practices for cultivating student scientific literacy. These difficulties may be overcome with newly available technologies, like generative AI systems, that enable personalized learning through domain-specific semantic spaces.

This article examines the instructional value of one such domain-specific LLM, Biodesign Buddy, through a mixed-methods analysis of an eight-week study involving 64 students participating in an international biodesign competition. Results indicate strong support for integrating AI into biodesign coursework. Surveys captured attitudes toward AI, scientific literature, and learning experiences to assess AI’s impact on learning outcomes. Findings suggest that integrating AI into biodesign pedagogy can meaningfully redress conceptual issues in biodesign while informing broader debates on AI’s role in higher education.

**Impact Statement:** This article introduces Biodesign Buddy, a domain-specific generative AI system for collegiate biodesign education, and reports on its exploratory deployment, offering design principles and preliminary findings to inform the development of AI-supported pedagogies for interdisciplinary biodesign instruction.

## Introduction

**Figure 1.**
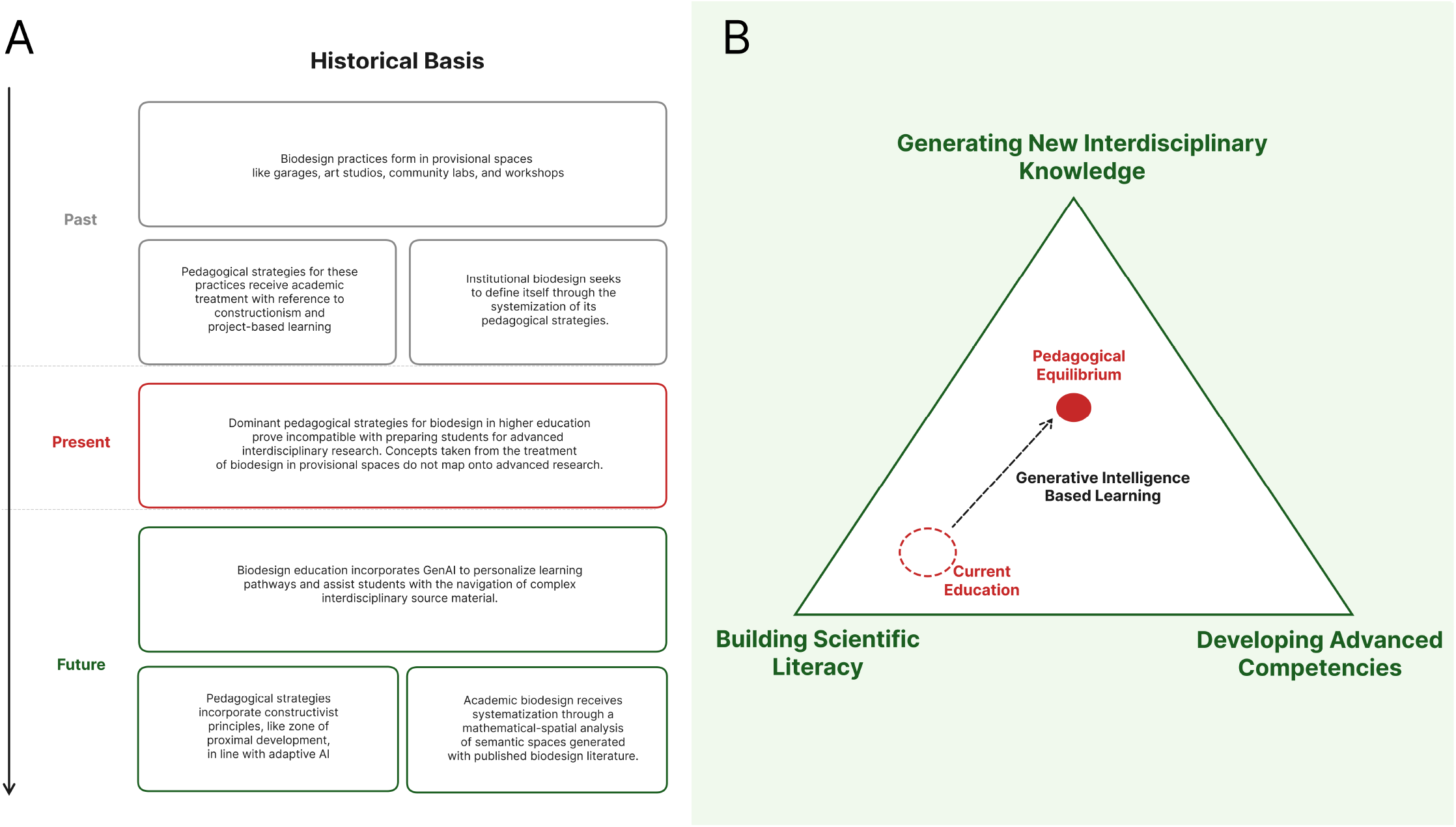
Conceptual framework for generative AI in biodesign pedagogy. (A) Historical trajectory from provisional making spaces through constructionist pedagogies to a GenAI-supported future. (B) Pedagogical triangle mapping current biodesign education against three competency dimensions, with Generative Intelligence Based Learning providing a pathway toward Pedagogical Equilibrium.

In a world of complex systems, hybrid academic studies have taken the place of traditional disciplines to better prepare students for emergent global challenges and an evolving workforce. This reorganization of higher education’s disciplinary structure marks a profound departure from the hierarchical, rule-bound orientation of conventional studies in an effort to better reflect the logics of contemporary problem solving (Newell and Klein, 1996). As a condition of this educational restructuring, integrative approaches to the natural sciences, like STEM and STEAM, have garnered particular attention for their ability to cultivate students’ intellectual curiosity, interdisciplinary competencies, and creative thinking abilities (Carter et al., 2021).

Biodesign, which draws lessons from both biotechnology and design, holds special promise among the newly formed hybrid academic models. Its application of life sciences research to ideas and products that promote global sustainability appeals widely to students and researchers (Pollini and Ragnoli, 2024). This interest, combined with the field’s potential positive impact on the planet, justifies present efforts to generate comprehensive educational strategies to benefit students who will eventually apply biodesign principles outside of higher education.

To further improve biodesign education, then, the field’s leaders may extend the logic of disciplinary integration by making generative artificial intelligence central to biodesign’s pedagogical strategies. In doing so, the adoption of these technologies can support students by facilitating rapid design iteration and personalized learning pathways. The decision to restructure current practices of biodesign education around the potential affordances of artificial intelligence, however, must be evaluated against the known risks. The potentially harmful effects of artificial intelligence on human creativity, academic honesty, and learning outcomes are already well established in educational studies literature. This article considers both the risks and the advantages of embracing AI in biodesign education through the comprehensive analysis of a domain-specific generative artificial intelligence system used in collegiate biodesign education.

### Literature Review

Biodesign is an emerging form of interdisciplinary research that incorporates principles from design and the life sciences to develop novel systems, processes, and objects that make essential use of biological functions (Myers, 2012). It occupies an important role in contemporary conversations around technoscientific futures, highlighting the economic and environmental possibilities that follow from advancements in biotechnology (Ginsberg and Chieza, 2018). In doing so, it attracts practitioners from diverse methodological backgrounds, forming an intellectual community that is defined more by the self-identification of its members than by a direct appeal to biotechnology and its normative concerns (Grushkin, 2021).

In the context of this shifting disciplinarity, biodesign has evolved from a niche activity conducted outside of academia into a mode of interdisciplinarity recognized within higher education. Top universities now offer undergraduate coursework in “biodesign” topics, and learning resources exist for prospective biodesign students at multiple levels of academic attainment (Danies et al., 2022; Raman, 2021).

Notably, the acceptance of biodesign methods within higher education can be mapped through the growth of the Biodesign Challenge, an international student design competition that encourages innovation in biotechnology, arts, and design. Founded in 2016, the Biodesign Challenge program began with representation from nine U.S. colleges and universities and now sees annual participation from more than sixty global universities, colleges, and secondary schools (Biodesign Challenge, 2025).

Because academic biodesign discourse is explicitly heterogeneous, students engaged in its formal educational pathways, like the Biodesign Challenge, encounter obstacles consistent with other forms of interdisciplinary design education, including the need to comprehend and synthesize incompatible perspectives from one or more discourses (Self et al., 2019). This cross-disciplinary approach, combined with skills like critical thinking and metacognition, is considered crucial for “future-proofing” design education and preparing students for industry (Brosens et al., 2022). Despite a growing consensus around the importance of interdisciplinary design education, however, questions remain about how to best prepare students for productive interdisciplinary work within a nascent discourse like biodesign.

The notion that biodesign is still an emerging field and, as such, lacks established pedagogical strategies and frameworks for disseminating knowledge informs the most current scholarship on biodesign education (Vijayakumar et al., 2024). This state of affairs frustrates collegiate educators who seek practical advice on biodesign curriculum design and its interdisciplinary objectives. Several conferences have supported research to address this issue, indicating possible interpretive frameworks for biodesign courses and key challenges to implementing them (Biodesign Challenge, 2024). At present, however, the standardization and sophistication of biodesign pedagogy, as opposed to biodesign research, finds its fullest expression outside of higher education.

Comprehensive pedagogical studies exist for biodesign instruction in both informal learning spaces and pre-collegiate contexts (Walker et al., 2022; Walker et al., 2023). These studies position biodesign as an alternative form of life sciences education with its emphasis on active learning, creativity, democratic access, and critical reflection (Walker et al., 2023). Such analyses draw an explicit connection between biodesign’s hands-on approach to life sciences learning and constructionism, an epistemological paradigm that emphasizes the role of public artefacts in knowledge formation (Papert and Harel, 1991; Kafai and Walker, 2020). The connection between constructionism and design thinking is well established (Kafai and Resnick, 2012; Holbert et al., 2020), making it an attractive paradigm for introductory biodesign investigations situated within the field’s conventional project-based learning scenarios. And yet, it may not prove suitable for advanced interdisciplinarity studies.

That constructionism, specifically, enjoys a privileged status in biodesign’s pedagogical thinking follows naturally from academic biodesign’s origins in DIY-biology and provisional spaces like kitchens, garages, and workshops (Pollini, 2024). Unlike formal scientific learning environments, which depend on specialized infrastructures and rigid empirical conventions, grassroot organizations like community laboratories support open-ended inquiry and free knowledge exchange (Jorgensen and Grushkin, 2011). These qualities can lead to the sort of interdisciplinary innovation typical of biodesign (Chappell et al., 2023). In such spaces, democratizing access to scientific discourse receives particular emphasis, and educational initiatives prioritize offering immediate engagement with biology with low barriers to entry. This arrangement differs from academic biodesign where, despite a shared commitment to democratizing biotechnology, the pedagogical priorities and schema of instruction must align with the institutional expectations of higher education.

Arguably, biodesign pedagogy at collegiate and post-collegiate levels should focus less on the acquisition of foundational biological concepts and more on the positive articulation of new interdisciplinary knowledge. To incentivize this intellectual production, educators might encourage biodesign students to undertake transdisciplinary material investigations (Marseglia et al., 2026), reflect on varieties of epistemic infrastructures (Ihls and Pollini, 2026), or work within more-than-human design philosophies (Zaretsky and Zaretsky, 2025). All have proven to be fruitful for encouraging student learning and creativity. However, for the students to fully participate in the proposed conversations, a minimum scientific literacy must be instilled through an approach to curriculum design that facilitates the assimilation and use of divergent, complex disciplinary perspectives.

Recently published research on student engagement strategies in biodesign points toward a viable pedagogical solution. A dual framework for teaching biodesign, which involves selectively adopting nature- or socially-driven approaches, has been demonstrated as a means for fostering interdisciplinary collaboration and improving scientific literacy (Danies et al., 2026). In this framework, biodesign educators appeal to the students’ interests and backgrounds to motivate them to learn unfamiliar methods in the service of biodesign innovation. This approach has been successfully explored in the context of the Biodesign Challenge, although not enough information is known about the students’ individual profiles and backgrounds to suggest general applicability across the field’s growing academic community.

Our article deviates from this approach by exploring an approach to biodesign education that incorporates generative artificial intelligence systems to assist students in navigating unfamiliar disciplinary frameworks. The potential contribution of these technologies, specifically large language models and domain-specific semantic spaces, has not yet been fully explored in the pedagogical literature on biodesign. One case study conducted in Colombia recently demonstrated that generative AI can assist undergraduate design students in the development of speculative biodesign projects (Ojeda Ramirez et al., 2026), but in doing so, it did not characterize how the technology could help cultivate the students’ scientific literacy for future design iterations. As one of the most comprehensive analysis of generative AI and biodesign education, however, two key topics from that study will inform the theoretical and empirical contributions of this article: speculative design and constructionist epistemology.

Revisiting these topics in light of the current initiative to reimagine biodesign pedagogies in reference to artificial intelligence comes at an important moment. The effort to formalize biodesign as a discipline through its pedagogical systematization coincides with generative artificial intelligence presenting substantive challenges to academia’s normative working conditions. Educational research has explored how higher education has already been affected by the adoption of these technologies in the classroom, particularly with respect to student assessment protocols, learning outcomes, and institutional policies (Shaw, 2025; Williams, 2025; McDonald et al., 2025).

The analysis of generative artificial intelligence in design education deserves special attention because art and design schools produce the most collegiate biodesign students. Studies note that, as with previous technological advancements, generative artificial intelligence introduces valuable new opportunities for the field while carrying substantive risk. The technical capabilities of generative AI lower barriers to entry, permitting the participation of new creative voices, while also raising questions about artistic authenticity and originality (Zailuddin et al., 2024). Equally, generative artificial intelligence can increase design student capacities, as in the case of sustainable design, to use complex interdisciplinary data sets (Li et al., 2025).

The effects of generative AI on natural sciences education appear more significant than they are in design education, partly due to more pervasive adoption (Karahan AdalI and Bigili, 2025). Analysis shows that, because generative artificial intelligence can outperform human operators in tasks that were formerly core to natural science disciplines, collegiate science education is undergoing substantive change. In response to the mathematical capacities of these technologies, emerging natural science pedagogies now place greater emphasis on cultivating diverse, interdisciplinary skillsets and student creativity (Tu et al., 2023).

The potential role of artificial intelligence in biodesign education will be addressed relative to these considerations and our findings in the discussion section of the article. Before doing so, however, we will discuss the technical design of Biodesign Buddy, the generative artificial intelligence systems given to students in the Biodesign Sprint competition, and the methods employed to study its impact on biodesign education.

### System Design and Implementation

#### Design Principles

The development of Biodesign Buddy was guided by three core design principles, each responding to a specific pedagogical challenge identified in the literature on interdisciplinary design education and AI-assisted learning

#### Transparency of analytical process

The first principle ensures that every output the system generates exposes its underlying reasoning. When Biodesign Buddy evaluates a student’s design concept, it does not return a score in isolation. Instead, it presents the analytical framework it applied, the dimensions it considered, the literature it retrieved, and the criteria it weighed. For instance, when the novelty assessment engine computes a similarity score between a student’s concept brief and existing scientific literature, the platform displays the specific PubMed articles it identified as analogs alongside a visual representation of their conceptual proximity. This transparency serves a dual pedagogical purpose: it models disciplined analytical thinking for students who may lack experience in systematic scientific evaluation, and it provides instructors with visibility into how the tool is shaping student reasoning.

#### Integration with External Evidence

The second principle grounds Biodesign Buddy’s outputs in verifiable scientific literature rather than relying solely on the language model’s parametric knowledge. The system queries PubMed via its public API to retrieve peer-reviewed research relevant to a student’s design hypothesis, embedding these references directly into its reports. This approach addresses a well-documented concern about large language models in educational contexts: their capacity to produce fluent but factually unsupported claims. By anchoring the tool’s guidance in externally retrievable sources, students can independently verify the scientific foundations of their projects, and instructors, who may themselves lack expertise in a specific subdomain of biology, can trust that the tool’s recommendations reflect published science rather than model confabulation.

#### Scaffolding Rather than Substitution

The third principle represents the most consequential design decision. Biodesign Buddy is explicitly designed to structure and support student inquiry, not to replace it. Rather than generating design concepts, writing project narratives, or producing finished analyses, the system provides frameworks, prompts, and reference materials that students must then interpret, synthesize, and apply through their own creative and analytical work. The platform’s guided flow, from concept brief to novelty report to learning pathway to iterative chat, is designed as a sequence of scaffolded interactions, each requiring active student engagement. This distinction reflects a constructionist learning philosophy: students learn most effectively when they are actively building something, and the tool’s role is to lower the barriers to that construction without performing it on the student’s behalf.

#### Technical Architecture

Biodesign Buddy is built on a decoupled architecture comprising a dynamic single-page web frontend and a serverless backend that handles all AI orchestration and data management. This separation of concerns ensures that the computational complexity of AI workflows remains isolated from the student-facing interface, producing a responsive user experience regardless of the processing demands of any given analytical task.

The backend is implemented as a set of cloud functions that serve as the secure orchestration layer for all AI-related workflows. Each analytical capability, novelty assessment, SWOT generation, learning pathway construction, and conversational chat, is implemented as a discrete, independently invocable function. This modular design means that each feature can be updated, tested, or scaled independently, and that the platform can expose only the capabilities appropriate to a given user’s role and project state. Role-based access control is enforced at the data layer through a multi-tenant security model distinguishing between global administrators, organization owners, and student members, ensuring that institutional data remains appropriately partitioned across schools and teams.

The AI orchestration layer uses a structured workflow framework to coordinate interactions between the underlying large language model and external data sources. All generative outputs are produced by a state-of-the-art multimodal language model accessed via a cloud AI platform, with model selection and versioning managed centrally to ensure consistency across student sessions. Critically, the orchestration layer is designed to interleave model inference with real-time retrieval from external sources rather than relying solely on the model’s parametric knowledge. This retrieval-augmented approach operationalizes the platform’s second design principle: by fetching live results from PubMed’s public API at query time, the system ensures that the scientific literature surfaced to students reflects the current state of published research rather than the model’s training distribution.

The novelty assessment pipeline illustrates this architecture in practice. When a student submits a concept brief, the system first encodes the text as a high-dimensional vector embedding using the language model’s representation layer. It then queries PubMed to retrieve a set of candidate articles whose abstracts are similarly encoded. Cosine similarity scores between the concept embedding and each retrieved abstract embedding are computed to produce a ranked analog set, which is visualized for the student as a proximity graph. The structured SWOT analysis is then generated by the language model with the retrieved articles and similarity scores passed as explicit context, grounding the model’s analytical outputs in the specific literature rather than in general domain knowledge. All project data, concept briefs, generated reports, chat histories, and learning pathways, persist in a document-oriented cloud database, enabling the conversational interface to maintain full project context across sessions without requiring students to re-establish their research state at each interaction.

#### Core Capabilities

**Figure 2.**
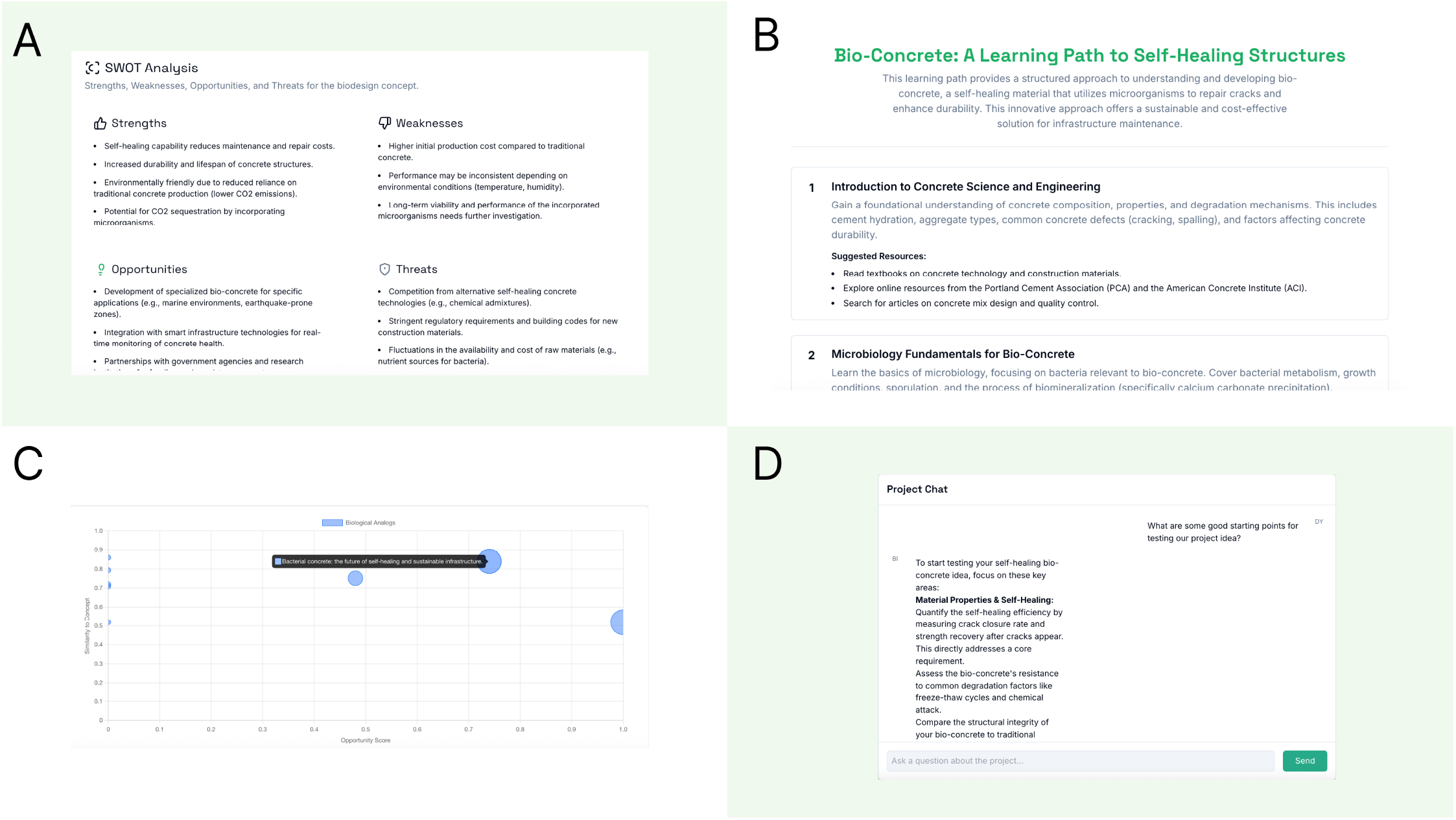
Biodesign Buddy interface displaying four core platform capabilities. (A) SWOT analysis generated from a student’s concept brief on bio-concrete, identifying project-specific strengths, weaknesses, opportunities, and threats grounded in retrieved scientific literature. (B) Personalized learning pathway structured as a sequenced curriculum of foundational and specialized knowledge resources tailored to the student’s project parameters. (C) Analog graph visualizing cosine similarity scores between the student’s concept and retrieved biological analogs, enabling students to locate their idea within the existing scientific landscape. (D) Context-aware project chat interface, retaining the concept brief and generated reports as persistent context to support iterative design refinement through open-ended dialogue.

Biodesign Buddy follows a linear, guided pedagogical flow designed to move students from an initial idea toward a structured research plan. This flow is organized around five interconnected capabilities.

The process begins with project initiation, where students create a new project by providing a title and a concept brief, a one-paragraph description of their bio-inspired design idea. This brief becomes the seed document from which all subsequent analyses are generated, and it anchors the AI’s context throughout the project lifecycle.

From the concept brief, students can generate a novelty assessment, which the platform terms a “Novelty Report.” This report integrates several analytical components. A SWOT analysis identifies project-specific strengths, weaknesses, opportunities, and threats, drawing on both the scientific literature and the design parameters articulated in the brief. Simultaneously, the system identifies real biological analogs, existing published research that is conceptually proximate to the student’s idea and presents these on an analog graph that visualizes the cosine similarity between the student’s concept and existing work. This visualization enables students to see where their idea sits relative to the scientific landscape, helping them identify both precedents and “white space” for innovation. For students from design backgrounds who may have limited experience with formal analytical frameworks, the structured SWOT output provides a disciplined entry point into systematic evaluation.

The learning pathways feature generates a personalized, ordered curriculum of skills and resources tailored to the specific technical requirements of the student’s project. Rather than presenting an undifferentiated list of papers, the system organizes relevant knowledge into a logical progression, from foundational concepts through specialized applications to specific technical skills (e.g., “Read articles on CRISPR,” “Learn Fusion 360”). This sequencing proved particularly valuable in the compressed eight-week sprint timeline, where students needed to build scientific understanding efficiently alongside their design work.

The conversational interface provides a context-aware project chat where the AI retains the project brief, and all generated reports as persistent context. This allows students to brainstorm follow-up questions, interrogate the SWOT analysis, explore the biology behind identified analogs, or refine their design concept through iterative dialogue, without re-explaining their project each time. Unlike the structured analytical tools, the chat supports open-ended exploration and sustained intellectual exchange.

Finally, a project management layer provides instructors and administrators with visibility into student research trajectories. Organization owners can manage team membership and access, while a global administrative interface allows oversight of user accounts and organization statuses. For instructors who are trained specialists in only one of biodesign’s constituent disciplines, this visibility is valuable: it enables them to monitor how teams are engaging with scientific literature and analytical tools, identify when students are struggling with unfamiliar territory, and intervene with targeted guidance even when the specific subject matter falls outside their own expertise.

**Figure 3.**
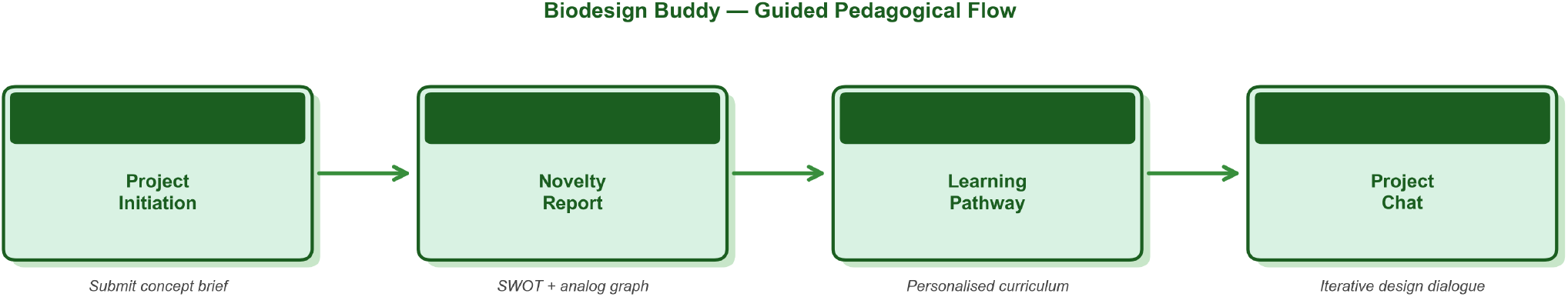
Biodesign Buddy Guided Pedagogical Flow Students progress from (1) project initiation through submission of a concept brief, to (2) generation of a literature-grounded novelty report incorporating SWOT analysis and analog mapping, to (3) construction of a personalized learning pathway, and finally to (4) iterative project dialogue via a context-aware conversational interface.

**Figure 4.**
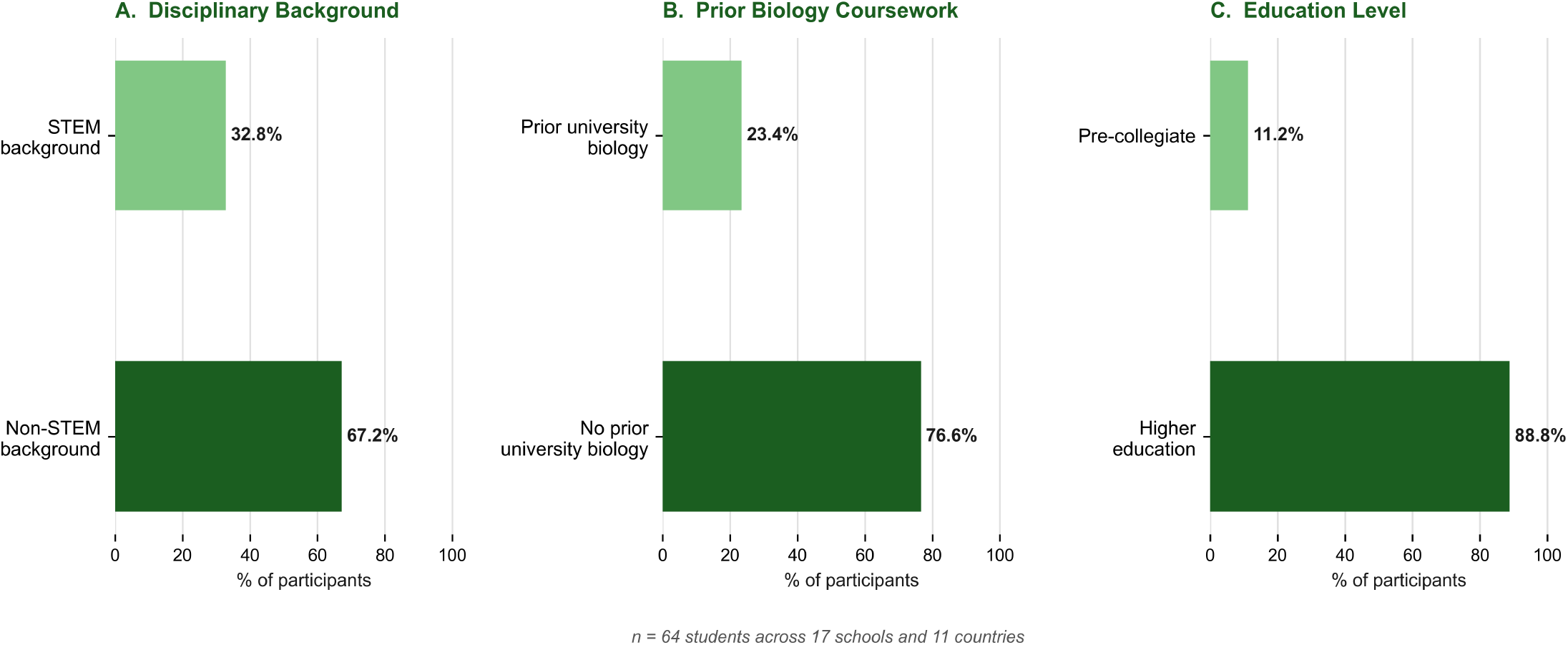
Participant Demographics and Educational Background. (A) Disciplinary background of students, showing a majority from non-STEM fields (67.2%) (B) Prior university-level biology coursework, with 76.6% reporting no formal biology training. (C) Education level, with most participants enrolled in higher education (88.8%) and a smaller proportion in pre-collegiate contexts (11.2%).

## Methods

### Deployment Context

Biodesign Buddy was deployed in the Biodesign Sprint: Ocean Futures program, an eight-week design competition hosted by Biodesign Challenge between October and December 2025. The Biodesign Sprint asked participating students to develop novel biodesign projects that aligned with the United Nations’ Sustainable Development Goals and its Ocean Decade framework. Educators enrolled their students in the Biodesign Sprint through their academic institutions, modifying the syllabi of preexisting art and design courses to comply with competition deliverables.

All educators and students associated with the Biodesign Sprint received supplemental academic support from the program organizers. Biodesign Challenge offered specialized webinars on topics related to ocean systems, access to a comprehensive research library, virtual design criticism sessions, and mentorship opportunities with biodesign experts. Access to Biodesign Buddy was provided as an optional, opt-in tool alongside a dedicated training webinar to familiarize students with the system’s capabilities and intended use.

A total of eighteen student teams, comprising 64 students, participated in the Biodesign Sprint. The student composition reflected diverse educational and cultural backgrounds, with representatives from seventeen schools and eleven countries submitting final projects for evaluation. The majority of students came from non-STEM backgrounds (67.2%), and a greater portion (76.6%) had no prior university-level biology coursework. Almost all teams (88.8%) were involved in higher education, with the remainder (11.2%) consisting of pre-collegiate learners. Due to the geographic distribution of participants, all program activities were conducted exclusively over Zoom.

To facilitate project evaluation, Biodesign Sprint organizers requested that each student team develop a ten-minute presentation underscoring the project’s unique contribution to biodesign, how scientific literature informed its development, and the problems it seeks to address. Like the mainstay Biodesign Challenge, evaluation of student projects relied on feedback from invited judges using a pre-distributed rubric divided into four core sections: narrative, concept, context, and reflection. Judges were not informed which teams had used Biodesign Buddy in the development of their projects.

### Data Collection

Biodesign Challenge and OpenTechBio organizers collected data throughout the program to assess the role of Biodesign Buddy in shaping student learning outcomes. Study organizers employed mixed-methods analysis on information collected directly from program participants through two surveys, as well as indirectly from the Sprint judges’ evaluations and submitted design notebooks.

The survey distributed prior to the Biodesign Sprint established baseline measurements for future analysis and collected demographic information about program participants. Respondents were asked questions pertaining to their educational background, including major discipline, as well as prior exposure to biodesign, self-reported scientific literacy, and highest level of academic achievement. Questions addressing beliefs about the importance of science in design education, human creativity, and the impact of artificial intelligence on the future of work were structured according to a five-point Likert scale.

A second survey distributed at the end of the program measured changes in student perceptions and beliefs over the course of the Biodesign Sprint. This exit survey revisited questions concerning student attitudes toward interdisciplinary research, science, artificial intelligence, and the future of the workforce. It also asked targeted questions about how access to Biodesign Buddy informed the design process and project formation. Among the 64 students who completed the pre-sprint survey, 13 completed the post-sprint survey, representing a response rate of 20.3%. Of those 13 respondents, 9 had access to Biodesign Buddy. Unless otherwise noted, all post-sprint quantitative findings are drawn from this respondent pool and should be interpreted as indicative rather than definitive.

An additional questionnaire was distributed to the instructors advising participating teams. This instrument captured demographic information, including educational attainment and disciplinary background, alongside Likert-scale items measuring attitudes toward science, artificial intelligence, and technology’s capacity to address complex social and environmental problems. Eighteen instructors completed this survey. Welch’s t-tests were used to identify statistically significant differences between student and instructor attitudes on shared items.

All participants with access to Biodesign Buddy were asked to produce design notebooks during the program. These notebooks were intended to provide records of interactions with the platform and its influence on the design process. Students received a template notebook outlining how to capture resources generated by Biodesign Buddy and reflect qualitatively on the iterative design process. The inclusion of written reflections on how AI-generated outputs shaped decision-making was explicitly encouraged. Design notebooks submitted at the program’s conclusion provided the primary source of qualitative evidence for this study.

## Results

Results are organized across five areas: adoption and usage patterns, perceived utility, qualitative feedback from design notebooks, challenges and limitations, and instructor perspectives. Unless otherwise noted, post-sprint survey data reflects a respondent pool of 13 students (20.3% response rate), of whom 9 had access to the Biodesign Buddy platform.

**Table 1.** Summary of quantitative findings across five measurement domains. Results are organized by (A) perceived utility of Biodesign Buddy among platform users (n=9), (B) engagement with scientific literature, (C) attitudinal shifts among all post-sprint respondents (n=13), (D) student versus instructor comparisons on shared Likert items assessed via Welch’s t-tests, and (E) instructor endorsement of AI in educational contexts (n=18). NA’s indicate measures collected at a single time point without a baseline comparator.

| Measure | Value / Pre-sprint | Post-sprint / Comparator |
| --- | --- | --- |
| **A. PERCEIVED UTILITY (post-sprint, platform users, n=9)** |  |  |
| Helpfulness: scientific foundations (mean / 5) | NA | 4.00 (SD 1.00) |
| - Non-STEM students (n=6) | NA | 4.17 |
| - STEM students (n=3) | NA | 3.67 |
| Helpfulness: chat functionality (mean / 5) | NA | 3.56 (SD 0.88) |
| Rated tool $\geq 4$ for scientific foundations | NA | 77.8% |
| **B. ENGAGEMENT WITH SCIENTIFIC LITERATURE** |  |  |
| Increased time on scientific literature (returning students) | NA | 75% (3 of 4) |
| Used other AI tools alongside platform | NA | 100% |
| **C. STUDENT ATTITUDINAL SHIFTS (all post-sprint, n=13)** |  |  |
| Designers should study science (1 - 5) | 4.19 | 4.46 |
| AI substitutes for human creativity (1 - 5) | 2.53 | 2.31 |
| **D. STUDENT vs. INSTRUCTOR COMPARISONS (Welch's t-tests)** |  |  |
| Comfort reading scientific papers (1 - 5) | Students: 3.61 | Instructors: 4.44 |
| Scientific literacy is important (1-5) | Students: 4.27 | Instructors: 4.67 |
| Designers should study science (1 - 5) | Students: 4.19 | Instructors: 4.56 |
| AI impact on traditional jobs (1 - 5) | Students: 4.02 | Instructors: 4.50 |
| AI substitutes for human creativity (1 - 5) | Students: 2.53 | Instructors: 2.61 |
| **E. INSTRUCTOR ENDORSEMENT OF AI (n=18)** |  |  |
| Encourage students to use AI tools | NA | 94.4% |
| Believe AI effective in classroom | NA | 88.9% |
| AI's effect on academic quality (mean / 5) | NA | 3.11 (SD 1.23) |

### Adoption and Usage Patterns

Of the 64 students who completed the pre-sprint survey, 98.4% (n = 63) reported prior experience with generative AI chatbots, and 75.0% (n = 48) described using them on a daily or weekly basis. The most reported AI functions were research (81.2%), image generation (46.9%), and chat-based advice and support (42.2%). This high baseline AI literacy meant that students entered the sprint with established expectations around AI tool use, providing important context for understanding their engagement with a domain-specific platform.

Among the 13 post-sprint respondents, 9 (69.2%) reported having access to Biodesign Buddy. The dominant usage pattern was iterative design refinement through the chat function, cited by 7 of 9 users. Fewer students reported using the tool primarily for conceptual organization and team workflow management (n = 2) or bibliography building (n = 2). When asked to describe their workflow, the most common sequence was project formation followed by literature search and chat interaction (n = 4), while 3 students described a more extended pipeline that included a subsequent project revision stage. Two students engaged only with the project formation and literature search features without using the chat function.

Notably, 100% of post-sprint respondents (n = 13) reported using other AI tools alongside Biodesign Buddy. ChatGPT was the most commonly cited complementary tool (n = 9), followed by Claude (n = 3), Gemini (n = 3), Microsoft Copilot (n = 2), and Perplexity (n = 3). This widespread co-use suggests that students treated Biodesign Buddy as a specialized complement to general-purpose AI platforms rather than a replacement for them, a pattern consistent with the tool’s design as a domain-specific scaffold rather than an all-purpose assistant.

### Perceived Utility

#### Scientific foundations

Students who had access to Biodesign Buddy rated its helpfulness in building a scientific foundation for their projects at a mean of 4.00 out of 5 (SD = 1.00, n = 9). Over three-quarters of users (77.8%) rated the tool at 4 or above, and a third (33.3%) gave it the highest possible rating. These results are notable given the disciplinary composition of the sample: 67.2% of pre-sprint participants came from non-STEM backgrounds, and 76.6% had never completed a college-level biology course. A comparison by disciplinary background reveals that non-STEM students rated Biodesign Buddy’s contribution to scientific foundations more highly (M = 4.17, n = 6) than their STEM counterparts (M = 3.67, n = 3). While the small sample sizes preclude inferential testing, this pattern is consistent with the tool’s design intent: students with less scientific preparation had more to gain from structured guidance through unfamiliar research domains, and their higher ratings suggest the scaffolding approach was effective in bridging that gap.

#### Chat functionality

Six of the 9 students with access (66.7%) used Biodesign Buddy’s chat functionality to refine their design concepts and scientific parameters. The mean usefulness rating for the chat was 3.56 out of 5 (SD = 0.88). While lower than the scientific foundations rating, this result likely reflects a more selective engagement pattern: students who used the chat tended to integrate it deeply into their iterative design process, while those who did not may have relied on general-purpose AI tools like ChatGPT for open-ended conversational interaction, a reasonable choice given the high rate of complementary tool use reported across the sample.

#### Time Efficiency and Engagement with Scientific Literature

The relationship between Biodesign Buddy and time spent navigating scientific literature produced a nuanced finding. Among the 4 returning students who could compare their experience to past biodesign projects, 3 reported that Biodesign Buddy increased the time they spent on scientific literature, while 1 reported a decrease. The remaining 5 respondents were participating in their first biodesign project and could not offer a comparison. Rather than indicating inefficiency, the increased time reported by most returning students suggests that the tool facilitated deeper engagement with scientific sources, directing students toward relevant research they might not have otherwise encountered. This interpretation is consistent with the platform’s scaffolding design principle, where the goal is to enrich the research process rather than abbreviate it. Overall, 75% of returning participants reported increased engagement with scientific literature over the course of the sprint.

### Attitudinal Shifts

Comparison of pre- and post-sprint survey responses among the 13 exit respondents revealed several noteworthy attitudinal shifts. Post-sprint responses showed greater endorsement of the view that designers should study science (M = 4.46, up from M = 4.19 at baseline), alongside a decreased belief that AI can substitute for human creativity (M = 2.31, down from M = 2.53). These directional changes, while modest in magnitude, suggest that sustained engagement with a science-anchored AI tool may reinforce rather than undermine students’ appreciation of human judgment and scientific grounding in design practice.

### Qualitative Feedback from Design Notebooks

Student design notebooks provided richer accounts of how Biodesign Buddy shaped project development. Two teams’ reflections are highlighted here for the depth and specificity of their assessments.

The University of Florence team described Biodesign Buddy’s contribution in terms of strategic orientation as much as informational support. In their reflection, they noted:

> *“The contribution provided by the Biodesign Buddy proved to be immediately valuable for acquiring a solid initial orientation, acting as a catalyst for the project’s launch. Furthermore, the accurate generation of the learning path constituted an indispensable strategic support, not only for facilitating the research setup phase but also for ensuring its structured and methodical progression throughout its entire development cycle*.*”*

This framing of the tool as a “catalyst” and a source of “strategic support” captures an important distinction between informational assistance and structural scaffolding. The Florence team did not describe the tool as telling them what to do but rather as providing a methodological orientation from which they could proceed with their own inquiry. This is precisely the register that the scaffolding design principle was intended to produce.

On the analog discovery tool, the team wrote:

> *“This tool provided some really interesting insight into some more research avenues for us - much more aligned with the ‘scientific’ paths of our research - most notably the chemical breakdowns of potential bio-technical material futures and the application of chemical compounds within our material recipe making and testing processes*.*”*

This observation confirms the analog discovery feature’s intended function: it did not simply return general information about biomaterials but surfaced specific scientific literature on chemical compound applications that became directly operative in the team’s material development process. The team went on to describe how the learning path feature structured their knowledge-building:

> *“This step allowed us to ensure that our base contextual knowledge for design development and the presentation is to define what the issue is, and hence understanding how we can begin to design solutions*.*”*

Critically, the Tasmania team’s reflection also documents a concrete design outcome that appears traceable to the tool’s research scaffolding. In their material testing process:

> *“A transition from material making to actual testing in sea water acted as a hurdle we had to face — chemically our recipes were supposed to last for 12-18 months but started degrading within minutes. This led to a swift recipe change which allowed for a greater end product*.*”*

This passage is significant because it illustrates the pathway from AI-guided literature engagement to real experimental consequences. The team’s ability to identify and respond to the chemical degradation problem was predicated on a research foundation that Biodesign Buddy helped construct. The team’s reflection represents one of the study’s clearest demonstrations that the platform’s scientific scaffolding had tangible effects on design outcomes.

The Tasmania team also identified one point of friction. On the learning path’s suggested prototyping stage, the team noted:

> *“This was probably the only learning path stage we felt was out of place, as we benefited from prototyping and testing much earlier in the process. As architecture students, I think we appreciate the practice of making and doing, and perhaps that is why we started the testing process much earlier*.*”*

This observation, combined with the team’s overall positive assessment, also yielded a concrete platform suggestion:

> *“A potential extension to the software is maybe a specific design journal, as a separate tab, where we can record our tests and prototypes outside of the AI chat section*.*”*

That a student team produced an actionable design critique of the platform they were using is itself an indication of the quality of engagement the tool elicited. The Tasmania team treated Biodesign Buddy as a research infrastructure worth improving, not merely a service to consume.

It is worth noting that the University of Tasmania received recognition in two award categories at the conclusion of the Biodesign Sprint, suggesting that their deep and critical engagement with the platform was consistent with strong project outcomes by external evaluation standards.

### Challenges and Limitations Identified

Several limitations emerged from the study. The most significant methodological constraint is the post-sprint response rate: only 13 of 64 participants (20.3%) completed the final survey, introducing potential non-response bias. Respondents may have been disproportionately engaged or satisfied with the program, and the small sample size limits the generalizability of quantitative findings. All statistical results should therefore be interpreted as indicative rather than definitive, and findings should be understood as generating hypotheses for future investigation rather than confirming them.

Student attrition during the eight-week sprint also affected the study. The compressed timeline and international, remote format presented logistical challenges for sustained participation, and not all teams who began the program completed final submissions. This pattern is consistent with broader challenges of maintaining engagement in remote, time-limited educational programs operating across multiple time zones.

### Instructor Perspectives

Eighteen instructors completed the pre-sprint survey, representing a range of disciplinary backgrounds: 44.4% (n = 8) in design fields, 38.9% (n = 7) in biological sciences, and 16.7% (n = 3) in environmental or sustainability studies. Over half (55.6%, n = 10) held degrees in the natural sciences, and the majority held doctoral (44.4%) or postdoctoral (16.7%) qualifications. This distribution reflects the interdisciplinary character of biodesign education, where most instructors are trained specialists in a single constituent discipline.

Instructor attitudes toward AI in education were overwhelmingly positive. Nearly all (94.4%, n = 17) reported encouraging students to use AI tools in their work, and 88.9% (n = 16) believed AI could be used effectively in the classroom. This high rate of endorsement is notable given that moderate concerns about privacy and intellectual property were also present (M = 3.56 out of 5, SD = 1.34), and that instructors rated AI’s effect on student academic quality at a modest 3.11 (SD = 1.23). This pattern suggests instructors recognized AI’s value as an educational tool while maintaining measured expectations about its direct impact on academic performance, a nuance that echoes the scaffolding-over-substitution principle embedded in the platform’s design.

Welch’s t-tests indicated statistically significant differences between student and instructor attitudes on four of six shared Likert items. Instructors reported significantly higher comfort reading scientific papers (M = 4.44 vs. 3.61, t = −3.43, p = .002), stronger endorsement of scientific literacy’s importance (M = 4.67 vs. 4.27, t = −2.35, p = .025), greater belief that designers should study science (M = 4.56 vs. 4.19, t = −2.15, p = .040), and higher assessment of AI’s impact on traditional jobs (M = 4.50 vs. 4.02, t = −2.29, p = .030). Notably, both groups scored similarly low on the belief that AI can substitute for human creativity (Students: M = 2.53; Instructors: M = 2.61, p = .792), indicating a shared skepticism about AI replacing the creative process.

## Discussion

Generative artificial intelligence systems, like Biodesign Buddy, offer biodesign students and educators an opportunity to enrich their interdisciplinary investigations at the intersection of biotechnology and design. With increased processing power and improved access to information, students who utilize these tools will find it easier to navigate interdisciplinary literature, test design hypotheses, and demonstrate creativity in the process of developing original ideas. Although this article explored the consequences of extending one generative artificial intelligence system to a modest number of users, the initial findings justify further case studies and discussion.

In conducting secondary studies, it is important to acknowledge that making generative artificial intelligence central to biodesign pedagogies will profoundly reshape the discipline. As discussed, pedagogical systematization plays an important role in formalizing biodesign as a distinct field in higher education. While biodesign’s academic identity remains underdetermined, the proposed centralization of artificial intelligence in biodesign pedagogy allows for the field to make important advances in the absence of a homogenous methodological framework or a shared set of concerns.

Importantly, citations, co-authorship, and other professional metrics can be encoded and mapped in a domain-specific semantic space for biodesign instruction, obviating the need for a scholarly consensus on a reductivist definition of biodesign. Such efforts are already underway, including one open-source LLM that biodesign researchers can currently use (SemanticSpace.ai, 2026). The adoption of this approach holds incredible potential to realign biodesign pedagogies with the standards of interdisciplinary practice exhibited in its academic journals. It may also, however, cause near-term confusion for students and educators accustomed to instructional models based on early biodesign pedagogies.

This proposed integration of generative artificial intelligence will have far-reaching effects in its challenge to two current orthodoxies of biodesign pedagogy: speculative design and constructionist epistemology. Both frameworks informed the study conducted with ChatGPT in Colombia but, as we will see, are complicated by student interactions with more robust generative intelligence systems.

Speculative design has been a mainstay of biodesign education. In the early years of academic biodesign, design students were encouraged to pursue speculative design projects due to the complexity of engaging with a specialized discipline like biotechnology (Pollini, 2024). In such cases, the adoption of speculative design pedagogies could be understood as a provisional use of a method, somewhat independent of its instructional value as a vehicle for interrogating the social and ethical dimensions of design (Dunne and Raby, 2013). For the advancement of the field, however, speculative design cannot continue in this function. Biodesign educators must carefully delineate between student projects that are speculative *by design* and those that are *speculative* due to limiting factors like inadequate resources (i.e. time, funds, and facilities) or a lack of scientific literacy. Generative artificial intelligence systems, like Biodesign Buddy, can assist in this process, providing responsive feedback on the validity and viability of student research questions. This clarity will improve academic assessment in the classroom by ensuring that ideas are appropriately categorized and motivate others to be carried further through prototyping and testing phases.

The challenge generative artificial intelligence will pose to the predominance of constructionist models in biodesign education is arguably more substantial. The constructionist paradigm applies to a very particular mode of instruction focused on prototyping and experimentation. As biodesign adopts more advanced learning technologies, its educators must evaluate whether constructionism continues to accurately represent the cognitive work being done by students. Although the previous study on AI in biodesign education persuasively connected its particular use of ChatGPT to constructionism, most potential uses of generative artificial intelligence will not focus on the manipulation of a visual object. In this case, entertaining alternative paradigms and their associated concepts can enrich our understanding of student learning in biodesign when investigations are mediated by an artificial agent and a curated semantic space.

Here, constructivism, the framework on which constructionism is based, deserves special attention. In contrast to Papert’s emphasis on public artefacts, Piaget’s constructivism proposes that learners construct knowledge by adding information to prior schema based on past experiences (Alanazi, 2016). In the literature, principles from both constructionism and constructivism have been used to describe biodesign adjacent learning (Walker et al., 2023), suggesting the appropriateness of a dual approach when describing how biodesign students may engage with relevant materials mediated by AI.

Lev Semenovich Vygotsky’s concept of the zone of proximal development, for instance, is uniquely well-suited for describing adaptive educational technologies that depend on mathematically measured distance. Vygotsky’s concept describes “the distance between the actual development level [of a learner] as determined by independent problem solving and the level of potential development as determined through problem solving under adult guidance or in collaboration with a more capable peer” (Vygotsky, 1978). Future studies should take advantage of archival functions of generative AI systems to mathematically understand how computational guidance contributes to students’ comprehension of interdisciplinary perspectives.

Beyond these broad disciplinary implications, generative artificial intelligence will undoubtedly affect the didactic benchmarks that educators aim to satisfy in guiding students through biodesign coursework. Biodesign educators must anticipate these changes in future curriculum design.

In adaptive educational systems, for example, artificial intelligence can support students by generating custom guidance introduced at a pace tailored to their abilities (Ayeni et al., 2024). This capability promises to address biodesign education’s longstanding issues with student scientific literacy. However, this benefit to adoption must be weighed against foreseeable educational challenges, like doubts over the students’ ability to evaluate the soundness of AI-generated content delivered in a personalized learning pathway (Murtaza et al., 2022). Additional issues may arise in the evaluation of student projects, as the personalized nature of guidance may lead students to pursue research outside the stated expertise of biodesign educators. Future studies will need to determine pedagogical best practices for these occasions.

Adjacent to learning pathway personalization issues, biodesign must formulate policies to inform its conception of ethical uses of artificial intelligence in the classroom. As noted elsewhere in educational studies literature, artificial intelligence introduces novel forms of academic dishonesty, which in turn negatively affect students, administrators, and educators (Shaw, 2025). Understanding how such forms of academic misconduct, like using GPT-generated text in assignments, might impact student assessment will be difficult, especially as professional work increasingly demands the use of information produced or modified by artificial intelligence. The successful formulation of these policies will necessitate additional case studies to disclose the ethical challenges that emerge as students experiment with artificial intelligence while developing biodesign projects.

## Conclusion

Biodesign education is entering a promising new era. Although academic biodesign currently lacks robust, widely distributed pedagogies for cultivating advanced interdisciplinary student research, emergent technologies, like generative artificial intelligence systems, introduce opportunities to reimagine the nature and depth of biodesign instruction. A limited study of one such system, Biodesign Buddy, suggests that generative artificial intelligence can support student creativity while fostering improved scientific literacy. Initial findings recommend further analyses and case studies in the context of educational competitions, like Biodesign Challenge, and a rethinking of fundamental ideas in biodesign pedagogical thought. In the future, generative artificial intelligence systems may become essential to effective biodesign instruction, empowering students to make novel contributions to the field.

## Funding Statement

This work was partially supported by unrestricted research funds awarded through the University of California, Irvine New Venture Competition. The funders had no role in study design, data collection, analysis, interpretation, or publication decisions.

## Competing Interests

Conflicts of Interest: Paul Rubery is employed at Biodesign Challenge. Dylan Riffle is the co-founder of OpenTechBio, the organization that developed Biodesign Buddy.

## Data Availability Statement

De-identified survey instruments and aggregated results are available on request from the corresponding author.

## Ethical Standards

This study involved minimal-risk educational research. All participants were informed about the study purpose and provided informed consent prior to participation. Participation was voluntary and participants could decline or withdraw at any time without penalty.

## Author Contributions

Conceptualization: D.R; P.R. Methodology: D.R; P.R. Data curation: D.R. Data visualisation: D.R. Writing original draft: D.R; P.R. All authors approved the final submitted draft.

## Notes

### Competing Interest Statement

Paul Rubery is employed by the Biodesign Challenge organization, which hosted the Biodesign Sprint program in which the study was conducted.
Dylan Riffle is the co-founder of OpenTechBio, the organization that developed the Biodesign Buddy platform evaluated in this study.

