## Supplementary File 1 for "Biodesign Buddy: Integrating Generative Artificial Intelligence in Academic Biodesign"

This document provides the complete survey instruments administered during the Biodesign Sprint: Ocean Futures study. Three instruments are included:

**S1** — Pre-Sprint Student Survey (n = 64): Baseline demographics, AI literacy, scientific background, and attitudinal measures.

**S2** — Post-Sprint Student Survey (n = 13; 20.3% response rate): Exit measures including attitudinal change items repeated from S1, and Biodesign Buddy-specific usage and utility items.

**S3** — Instructor Survey (n = 18): Pre-sprint demographics, disciplinary background, and attitudes toward AI and science education.

### S1. Pre-Sprint Student Survey

Administered at program outset to all enrolled students. Establishes baseline measures for demographic composition, prior AI and scientific experience, and attitudinal benchmarks used in pre-post comparisons (Table 1 of main manuscript).

#### Section A: Demographics and Educational Background

| **Q1** | **What is your current level of study?** |
|---|---|
| Type | *Select one* |
| Options | High School Student |
| | Undergraduate at College/University |
| | Masters Student |
| | Doctoral Student (PhD) |
| | Post-Doctoral Researcher |
| | Professional (non-student) |

| **Q2** | **Are you a native English speaker?** |
|---|---|
| Type | *Select one* |
| Options | Yes |
| | No |
| | Native Bilingual Proficiency |

| **Q3** | **How many years of experience do you have in biodesign and/or related topics?** |
|---|---|
| Type | *Select one* |
| Options | 1 year or less |
| | 1–2 years |
| | 3–5 years |

| | |
|---|---|
| | 5+ years |

| Q4 | Which of the following best describes your college major or course of study? |
|---|---|
| Type | *Select one* |
| Options | Studio Art, Design, or Related Discipline |
| | Biology, Life Sciences, or Related Discipline |
| | Engineering or Related Discipline |
| | Environmental or Sustainability Studies |
| | Social Sciences or Humanities |
| | Other |

| Q5 | If you are enrolled in a design school or program, select which field best describes your area of focus: |
|---|---|
| Type | *Select one* |
| Options | Architecture |
| | Industrial Design |
| | Graphic / Communication Design |
| | Fashion Design |
| | Interior Design |
| | N/A |
| | Other |

| Q6 | Which of the following best describes your current location? |
|---|---|
| Type | *Select one* |
| Options | North America |
| | South America |
| | Europe |
| | Asia |
| | Africa |
| | Oceania |
| | Middle East |

### Section B: Technology and AI Background

| Q7 | Do you regularly use any of the following Adobe products or their open-source competitors: Photoshop, Illustrator, InDesign, Premiere Pro, After Effects, Lightroom? |
|---|---|
| Type | *Select one* |
| Options | Yes |

| | |
|---|---|
| | No |

| Q8 | **Do you know how to use Blender?** |
|---|---|
| Type | *Select one* |
| Options | Yes |
| | No |

| Q9 | **Have you ever used a generative artificial intelligence chatbot like ChatGPT, DeepSeek, or Gronk?** |
|---|---|
| Type | *Select one* |
| Options | Yes |
| | No |

| Q10 | **If you have used generative artificial intelligence chatbots before, which of the following best describes how often you have used them?** |
|---|---|
| Type | *Select one* |
| Options | Daily |
| | Weekly |
| | Once or Twice a Month |
| | Rarely |
| | Never |

| Q11 | **Does your school and/or educational institution have a policy regulating the use of artificial intelligence (AI) in the classroom or on assignments?** |
|---|---|
| Type | *Select one* |
| Options | Yes |
| | No |

| Q12 | **Please tell us what functions you use general artificial intelligence for.** |
|---|---|
| Type | *Select all that apply* |
| Options | Research |
| | Image Generation |
| | Chat Functionality (Advice, support, companionship, etc.) |
| | Composing emails and other texts |
| | Entertainment |
| | Basic understanding of topics |
| | Other |

### Section C: Scientific Literacy and Laboratory Experience

| Q13 | Have you completed a college/university level course in biology or a related life sciences discipline? |
| --- | --- |
| Type | *Select one* |
| Options | Yes |
| | No |

| Q14 | Have you ever conducted an experiment in a laboratory? |
| --- | --- |
| Type | *Select one* |
| Options | Yes |
| | No |

| Q15 | Do you know how to structure, arrange, and keep a laboratory notebook? |
| --- | --- |
| Type | *Select one* |
| Options | Yes |
| | No |

| Q16 | How would you describe your comfort level with reading and understanding scientific research papers? |
| --- | --- |
| Type | *5-point scale* |
| Options | 1 — Very difficult to understand |
| | 2 |
| | 3 — Can understand general content |
| | 4 |
| | 5 — Very comfortable reading and analyzing |

### Section D: Attitudes Toward Science, Design, and AI

*Items Q17–Q21 were repeated in the post-sprint survey (S2) to measure attitudinal change over the program duration.*

| Q17 | How important do you think scientific literacy is to effectively navigate modern life? |
| --- | --- |
| Type | *5-point Likert scale (1 = Not at all important, 5 = Extremely important)* |
| Options | 1 — Strongly Disagree / Not at all |
| | 2 |
| | 3 — Neutral |
| | 4 |
| | 5 — Strongly Agree / Extremely |

| Q18 | How important do you think it is for designers to study scientific topics in school? |
| --- | --- |

| | |
|---|---|
| Type | *5-point Likert scale (1 = Not at all important, 5 = Extremely important)* |
| Options | 1 — Strongly Disagree / Not at all |
| | 2 |
| | 3 — Neutral |
| | 4 |
| | 5 — Strongly Agree / Extremely |

| **Q19** | **How probable do you think it is that advanced technology will solve environmental problems like climate change?** |
|---|---|
| Type | *5-point Likert scale (1 = Not at all probable, 5 = Extremely probable)* |
| Options | 1 — Strongly Disagree / Not at all |
| | 2 |
| | 3 — Neutral |
| | 4 |
| | 5 — Strongly Agree / Extremely |

| **Q20** | **To what extent do you think artificial intelligence will impact traditional jobs and occupations?** |
|---|---|
| Type | *5-point Likert scale (1 = Not at all, 5 = Extremely)* |
| Options | 1 — Strongly Disagree / Not at all |
| | 2 |
| | 3 — Neutral |
| | 4 |
| | 5 — Strongly Agree / Extremely |

| **Q21** | **How strongly do you believe that artificial intelligence can substitute for human creativity?** |
|---|---|
| Type | *5-point Likert scale (1 = Not at all, 5 = Completely)* |
| Options | 1 — Strongly Disagree / Not at all |
| | 2 |
| | 3 — Neutral |
| | 4 |
| | 5 — Strongly Agree / Extremely |

| **Q22** | **Are you optimistic about artificial intelligence and other future technologies?** |
|---|---|
| Type | *Select one* |
| Options | Yes |
| | No |
| | I don't know / unsure |

#### Section E: Program Goals and Expectations

| Q23 | What do you hope to accomplish in the Biodesign Sprint: Ocean Futures program? |
| --- | --- |
| Type | *Select all that apply* |
| Options | Develop a project for a design portfolio |
| | Promote my work to a wider audience |
| | Win a prize/receive recognition in an international competition |
| | Form professional relationships with the biodesign community |
| | Learn more about biodesign as an emerging design discipline |
| | Become more familiar with AI-assisted learning protocols |
| | Other |

| Q24 | How much do you think this competition will prepare you for the AI workforce of the future? |
| --- | --- |
| Type | *5-point Likert scale* |
| Options | 1 — Strongly Disagree / Not at all |
| | 2 |
| | 3 — Neutral |
| | 4 |
| | 5 — Strongly Agree / Extremely |

| Q25 | (Optional) Is there anything you would like to share with the study organizers before the program begins? |
| --- | --- |
| Type | *Open-ended text response* |

### S2. Post-Sprint Student Survey

Administered at program conclusion (n = 13; 20.3% response rate). Of these, 9 reported access to Biodesign Buddy. Section A repeats attitudinal items from S1 for pre-post comparison. Sections C–D items were conditionally displayed only to respondents who indicated platform access.

#### Section A: Repeated Attitudinal Items

*Scale anchors are identical to S1 Section D to enable pre-post comparison.*

| Q1 | What is your current level of study? |
| --- | --- |
| Type | *Select one* |
| Options | High School Student |
| | Undergraduate at College/University |
| | Masters Student |

| | |
|---|---|
| | Doctoral Student (PhD) |
| | Post-Doctoral Researcher |
| | Professional (non-student) |

| **Q2** | **How would you describe your comfort level with reading and understanding scientific research papers?** |
|---|---|
| Type | *5-point scale — repeated from pre-sprint survey* |
| Options | 1 — Very difficult to understand |
| | 2 |
| | 3 — Can understand general content |
| | 4 |
| | 5 — Very comfortable reading and analyzing |

| **Q3** | **How important do you think scientific literacy is to effectively navigate modern life?** |
|---|---|
| Type | *5-point Likert scale — repeated from pre-sprint survey for pre-post comparison* |
| Options | 1 — Strongly Disagree / Not at all |
| | 2 |
| | 3 — Neutral |
| | 4 |
| | 5 — Strongly Agree / Extremely |

| **Q4** | **How important do you think it is for designers to study scientific topics in school?** |
|---|---|
| Type | *5-point Likert scale — repeated from pre-sprint survey for pre-post comparison* |
| Options | 1 — Strongly Disagree / Not at all |
| | 2 |
| | 3 — Neutral |
| | 4 |
| | 5 — Strongly Agree / Extremely |

| **Q5** | **To what extent do you think artificial intelligence will impact traditional jobs and occupations?** |
|---|---|
| Type | *5-point Likert scale — repeated from pre-sprint survey for pre-post comparison* |
| Options | 1 — Strongly Disagree / Not at all |
| | 2 |
| | 3 — Neutral |
| | 4 |
| | 5 — Strongly Agree / Extremely |

| **Q6** | **How strongly do you believe that artificial intelligence can substitute for human creativity?** |
|---|---|

| | |
|---|---|
| Type | *5-point Likert scale — repeated from pre-sprint survey for pre-post comparison* |
| Options | 1 — Strongly Disagree / Not at all |
| | 2 |
| | 3 — Neutral |
| | 4 |
| | 5 — Strongly Agree / Extremely |

| **Q7** | **Can humans direct artificial intelligence to benefit society?** |
|---|---|
| Type | *Select one* |
| Options | Yes |
| | No |
| | I don't know / unsure |

| **Q8** | **Do you believe artificial intelligence can accelerate student learning of scientific concepts?** |
|---|---|
| Type | *Select one* |
| Options | Strongly agree |
| | Agree |
| | Neutral |
| | Disagree |
| | Strongly disagree |

| **Q9** | **Do you believe that artificial intelligence can be used by design students to supplement traditional areas of competency?** |
|---|---|
| Type | *Select one* |
| Options | Strongly agree |
| | Agree |
| | Neutral |
| | Disagree |
| | Strongly disagree |

| **Q10** | **Does artificial intelligence make it easier for scientists or designers to work in an interdisciplinary fashion?** |
|---|---|
| Type | *Select one* |
| Options | Strongly agree |
| | Agree |
| | Neutral |
| | Disagree |
| | Strongly disagree |

### Section B: AI Tool Use

| Q11 | Did you have access to the Biodesign Buddy tool during the Biodesign Sprint? |
|---|---|
| Type | *Select one* |
| Options | Yes |
| | No |

| Q12 | Select the answers that best describe how you used the Biodesign Buddy tool during the Biodesign Sprint: Ocean Futures program. |
|---|---|
| Type | *Select all that apply \| Displayed to respondents who indicated Biodesign Buddy access* |
| Options | Iterative Design: Interacting with the Chat Function to Refine Design Concepts |
| | Conceptual Organization: Managing Workflow in a Team Dynamic |
| | Bibliography Building: Using the Literature Search to Organize Academic Sources |
| | Other |

| Q13 | Did you use other AI tools to develop your final project? |
|---|---|
| Type | *Select one* |
| Options | Yes |
| | No |

| Q14 | If you used other AI tools, which tools did you use? |
|---|---|
| Type | *Select all that apply* |
| Options | ChatGPT |
| | Claude |
| | Gemini |
| | Microsoft Copilot |
| | Perplexity |
| | DeepSeek |
| | Other |

### Section C: Biodesign Buddy Perceived Utility

*Items Q15–Q22 were displayed only to respondents who indicated access to Biodesign Buddy (n = 9).*

| Q15 | On a sliding scale, how helpful was Biodesign Buddy in building a scientific foundation for your final project? |
|---|---|
| Type | *5-point sliding scale \| Displayed to respondents who indicated Biodesign Buddy access* |

| | |
|---|---|
| Options | 1 (Not at all helpful) |
| | 2 |
| | 3 |
| | 4 |
| | 5 (Extremely helpful) |

| **Q16** | **Did you use Biodesign Buddy's chat functionality to refine your design concept and its scientific parameters?** |
|---|---|
| Type | *Select one \| Displayed to respondents who indicated Biodesign Buddy access* |
| Options | Yes |
| | No |

| **Q17** | **On a sliding scale, how useful was Biodesign Buddy's chat functionality in testing and revising project ideas?** |
|---|---|
| Type | *5-point sliding scale \| Displayed to respondents who used chat functionality* |
| Options | 1 (Not at all useful) |
| | 2 |
| | 3 |
| | 4 |
| | 5 (Extremely useful) |

| **Q18** | **Which answer best describes your workflow with Biodesign Buddy?** |
|---|---|
| Type | *Select one \| Displayed to respondents who indicated Biodesign Buddy access* |
| Options | Project Formation → Literature Search |
| | Project Formation → Literature Search → Chat Function |
| | Project Formation → Literature Search → Chat Function → Project Revision |
| | Other |

| **Q19** | **Do you think your project would have changed dramatically without access to Biodesign Buddy?** |
|---|---|
| Type | *Select one \| Displayed to respondents who indicated Biodesign Buddy access* |
| Options | Yes |
| | No |
| | Maybe |

| **Q20** | **Did your team explore multiple project ideas with Biodesign Buddy?** |
|---|---|
| Type | *Select one \| Displayed to respondents who indicated Biodesign Buddy access* |
| Options | Yes |
| | No |

| | |
|---|---|
| | Maybe |

| **Q21** | **How did Biodesign Buddy affect the time you spent navigating scientific literature relative to past biodesign projects?** |
|---|---|
| Type | *Select one \| Displayed to respondents who indicated Biodesign Buddy access* |
| Options | Increased |
| | Decreased |
| | No change |
| | N/A: First Biodesign Project |

| **Q22** | **What functionality would you like to see added to Biodesign Buddy for future students and users?** |
|---|---|
| Type | *Open-ended text response \| Displayed to respondents who indicated Biodesign Buddy access* |

### Section D: Program Outcomes

| **Q23** | **What did you accomplish in the Biodesign Sprint: Ocean Futures program?** |
|---|---|
| Type | *Select all that apply* |
| Options | Develop a project for a design portfolio |
| | Promote your work to a wider audience |
| | Win a prize/receive recognition in an international competition |
| | Form professional relationships with the biodesign community |
| | Learn more about biodesign as an emerging design discipline |
| | Become more familiar with AI-assisted learning protocols |

| **Q24** | **How much do you think this competition prepared you for the AI workforce of the future?** |
|---|---|
| Type | *5-point Likert scale* |
| Options | 1 — Strongly Disagree / Not at all |
| | 2 |
| | 3 — Neutral |
| | 4 |
| | 5 — Strongly Agree / Extremely |

| **Q25** | **(Optional) Is there anything you would like to share with the study organizers about Biodesign Buddy and its use in the classroom?** |
|---|---|
| Type | *Open-ended text response* |

### S3. Instructor Survey

Administered to all instructors advising participating teams prior to program commencement (n = 18). Shared Likert items with the student survey (Q17–Q21) were used for Welch's t-test comparisons reported in Table 1D of the main manuscript.

#### Section A: Demographics and Professional Background

| **Q1** | **What is your highest level of educational attainment?** |
|---|---|
| Type | *Select one* |
| Options | Undergraduate Degree |
| | Masters Degree |
| | Doctoral Degree (PhD) |
| | Post-Doctoral |

| **Q2** | **What best describes your area of professional expertise?** |
|---|---|
| Type | *Select one* |
| Options | Studio Art, Design, or Related Discipline |
| | Biology, Life Sciences, or Related Discipline |
| | Engineering or Related Discipline |
| | Environmental or Sustainability Studies |
| | Social Sciences or Humanities |
| | Other |

| **Q3** | **How many years of experience do you have in biodesign and/or related topics?** |
|---|---|
| Type | *Select one* |
| Options | 1 year or less |
| | 1–2 years |
| | 3–5 years |
| | 5+ years |

| **Q4** | **What level of student are you advising/teaching in the Biodesign Sprint: Ocean Futures program?** |
|---|---|
| Type | *Select one* |
| Options | High School |
| | Undergraduate |
| | Masters |
| | Mixed levels |

| Q5 | Are you a native English speaker? |
|---|---|
| Type | *Select one* |
| Options | Yes |
| | No |
| | Native Bilingual Proficiency |

| Q6 | Which of the following best describes your location? |
|---|---|
| Type | *Select one* |
| Options | North America |
| | South America |
| | Europe |
| | Asia |
| | Africa |
| | Oceania |
| | Middle East |

### Section B: Technology and AI Background

| Q7 | Do you regularly use any of the following Adobe products or their open-source competitors? |
|---|---|
| Type | *Select one* |
| Options | Yes |
| | No |

| Q8 | Have you ever used a generative artificial intelligence chatbot like ChatGPT, DeepSeek, or Gronk? |
|---|---|
| Type | *Select one* |
| Options | Yes |
| | No |

| Q9 | How often do you use generative AI chatbots? |
|---|---|
| Type | *Select one* |
| Options | Daily |
| | Weekly |
| | Once or Twice a Month |
| | Rarely |
| | Never |

| Q10 | **Does your school and/or educational institution have a policy regulating the use of artificial intelligence (AI) in the classroom or on assignments?** |
|---|---|
| Type | *Select one* |
| Options | Yes |
| | No |

| Q11 | **Do you have a written policy regulating the use of artificial intelligence (AI) on assignments?** |
|---|---|
| Type | *Select one* |
| Options | Yes |
| | No |

| Q12 | **Please tell us what functions you use general artificial intelligence for.** |
|---|---|
| Type | *Select all that apply* |
| Options | Research |
| | Image Generation |
| | Chat Functionality (Advice, support, companionship, etc.) |
| | Composing emails and other texts |
| | Entertainment |
| | Basic understanding of topics |
| | Other |

### Section C: Scientific Background

| Q13 | **Do you possess a degree in the natural sciences?** |
|---|---|
| Type | *Select one* |
| Options | Yes |
| | No |

| Q14 | **Have you ever conducted an experiment in a laboratory?** |
|---|---|
| Type | *Select one* |
| Options | Yes |
| | No |

| Q15 | **Do you know how to structure, arrange, and keep a laboratory notebook?** |
|---|---|
| Type | *Select one* |
| Options | Yes |
| | No |

| Q16 | How would you describe your comfort level with reading and understanding scientific research papers? |
|---|---|
| Type | *5-point scale* |
| Options | 1 — Very difficult to understand |
| | 2 |
| | 3 — Can understand general content |
| | 4 |
| | 5 — Very comfortable reading and analyzing |

### Section D: Attitudes Toward Science, Design, and AI

*Items Q17–Q21 are shared with the student survey and were used for Welch's t-test comparisons between student and instructor groups.*

| Q17 | How important do you think scientific literacy is to effectively navigate modern life? |
|---|---|
| Type | *5-point Likert scale — shared item with student survey; used for Welch's t-test comparison* |
| Options | 1 — Strongly Disagree / Not at all |
| | 2 |
| | 3 — Neutral |
| | 4 |
| | 5 — Strongly Agree / Extremely |

| Q18 | How important do you think it is for designers to study scientific topics in school? |
|---|---|
| Type | *5-point Likert scale — shared item with student survey; used for Welch's t-test comparison* |
| Options | 1 — Strongly Disagree / Not at all |
| | 2 |
| | 3 — Neutral |
| | 4 |
| | 5 — Strongly Agree / Extremely |

| Q19 | How probable do you think it is that advanced technology will solve environmental problems like climate change? |
|---|---|
| Type | *5-point Likert scale* |
| Options | 1 — Strongly Disagree / Not at all |
| | 2 |
| | 3 — Neutral |
| | 4 |
| | 5 — Strongly Agree / Extremely |

| Q20 | To what extent do you think artificial intelligence will impact traditional jobs and occupations? |
|---|---|
| Type | *5-point Likert scale — shared item with student survey; used for Welch's t-test comparison* |
| Options | 1 — Strongly Disagree / Not at all |
| | 2 |
| | 3 — Neutral |
| | 4 |
| | 5 — Strongly Agree / Extremely |

| Q21 | How strongly do you believe that artificial intelligence can substitute for human creativity? |
|---|---|
| Type | *5-point Likert scale — shared item with student survey; used for Welch's t-test comparison* |
| Options | 1 — Strongly Disagree / Not at all |
| | 2 |
| | 3 — Neutral |
| | 4 |
| | 5 — Strongly Agree / Extremely |

| Q22 | To what extent do privacy and/or intellectual property concerns affect your attitudes towards artificial intelligence? |
|---|---|
| Type | *5-point Likert scale* |
| Options | 1 — Strongly Disagree / Not at all |
| | 2 |
| | 3 — Neutral |
| | 4 |
| | 5 — Strongly Agree / Extremely |

| Q23 | What effect do you think artificial intelligence has on the quality of your students' academic performance? |
|---|---|
| Type | *5-point scale* |
| Options | 1 (Very negative effect) |
| | 2 |
| | 3 (Neutral) |
| | 4 |
| | 5 (Very positive effect) |

| Q24 | Do you believe that artificial intelligence can be used effectively in the classroom? |
|---|---|
| Type | *Select one* |
| Options | Yes |

| | |
|---|---|
| | No |
| | Maybe / Unsure |

| **Q25** | **Are you optimistic about artificial intelligence and other future technologies?** |
|---|---|
| Type | *Select one* |
| Options | Yes |
| | No |
| | I don't know / unsure |

| **Q26** | **Do you currently encourage students to take advantage of artificial intelligence tools in their work?** |
|---|---|
| Type | *Select one* |
| Options | Yes |
| | No |

### Section E: Program Goals

| **Q27** | **What do you hope for your students to accomplish in the Biodesign Sprint?** |
|---|---|
| Type | *Select all that apply* |
| Options | Develop a project for a design portfolio |
| | Promote their work to a wider audience |
| | Win a prize/receive recognition in an international competition |
| | Form professional relationships within the biodesign community |
| | Learn more about biodesign as an emerging design discipline |
| | Become more familiar with AI-assisted learning protocols |

| **Q28** | **What do YOU hope to accomplish by leading students through the Biodesign Sprint?** |
|---|---|
| Type | *Select all that apply* |
| Options | Win a prize/receive recognition for the school |
| | Form professional relationships within the biodesign community |
| | Learn more about biodesign as an emerging design discipline |
| | Become more familiar with AI-assisted learning protocols |
| | Other |

| **Q29** | **(Optional) Is there anything you would like to share with the study organizers before the program begins?** |
|---|---|
| Type | *Open-ended text response* |
